# Are we overestimating the utility of hair glucocorticoids? A systematic review exploring the empirical evidence supporting hair glucocorticoids as a measure of stress

**DOI:** 10.1101/375667

**Authors:** Otto Kalliokoski, Finn K. Jellestad, Robert Murison

**Affiliations:** Department of Experimental Medicine, University of Copenhagen, Denmark; Department of Biological and Medical Psychology, University of Bergen, Norway

**Keywords:** Hair, Glucocorticoids, Cortisol, Corticosterone, Stress

## Abstract

Quantitating glucocorticoids (GCs) in hairs is a popular method for assessing chronic stress in studies of humans and animals alike. The cause-and-effect relationship between stress and elevated GC levels in hairs, sampled weeks later, is however hard to prove. This systematic review evaluated the evidence supporting hair glucocorticoids (hGCs) as a biomarker of stress.

Only a relatively small number of controlled studies employing hGC analyses have been published, and the quality of the evidence is compromised by unchecked sources of bias. Subjects exposed to stress mostly demonstrate elevated levels of hGCs, and these concentrations correlate significantly with GC concentrations in serum, saliva and feces. This supports hGCs as a biomarker of stress, but the dataset provided no evidence that hGCs are a marker of historical stress. Only in cases where the stressor persisted at the time of hair sampling could a clear link between stress and hGCs be established.

## 1. Background

Measuring glucocorticoids (GCs) deposited in hair is an increasingly popular method for biomarker-based stress assessment. Hair is sampled easily and painlessly, it is often an abundant source of material, and it has been argued to have superior qualities over other methods for analyzing GCs when it comes to gauging chronic stress^1–3^. If GCs are sequestered from the blood stream and locked into place in the growing hair at the level of the hair follicle, a single strand of hair contains within it a historical record of the HPA axis activity of its owner spanning months into the past. This idea is taken to the next level when researchers segment hairs and analyze different sections, ostensibly corresponding to different periods in the past, to make inferences regarding the perceived stress levels over time of their subjects, whether human patients^4^, captive animals^5^, or long-dead mummies^6^.

But are hair glucocorticoids (hGCs) a robust marker of stress? With local production of GCs in the hair follicle^7^ – the local HPA axis appearing to respond to local stressors independently of the rest of the organism^8^ – an uncertain rate of incorporation of GCs into hairs^9^ and unknown mechanisms by which this takes place^10^, it is unclear to which degree hGCs are reflective of the (central) stress response of an individual. Moreover, with some evidence that hGC concentrations change much faster than can be explained by mechanisms concerning only incorporation of GCs in the hair follicle^11,12^, can sections of a hair really be related to a specific period in the past?

Despite the many unknowns surrounding the use of hGCs as a measure of chronic stress, the biomarker is presently used to gauge mental illness^13^, the wellbeing of human trauma victims^14^ (and long-term consequences of the trauma^15,16^), post-traumatic stress disorder (PTSD) sufferers^4,17,18^, and children^19,20^; to assess animal welfare in wildlife^21^, captive animals^22,23^ and laboratory animals^24^. The mismatch between the uncertainties of the method and the confidence with which it is applied is concerning.

The present systematic review strived to collect and evaluate the empirical evidence supporting the use of hGC analyses as a method for assessing physiological and psychological stress. With the method having been originally developed for studying stress in wildlife^2^, but presently being frequently used to investigate human patients^3^, limiting the study to either humans or non-human animals would not have painted a complete picture. Unlike previous reviews/meta-analyses^3,13,25,26^, we thus set out to collate data from all mammalian species; to include studies carried out in human and non-human animals alike.

## 2. Material and methods

The methods listed below were pre-specified in a study protocol accessible online since Jan 13, 2016 (Supplemental materials A).

A broad – inclusive – search strategy was employed in an attempt to find all relevant publications that could provide unambiguous evidence of hGCs being related to the HPA-axis-activating stress response of an individual. The findings of the studies included in the present review were judged qualitatively as well as quantitatively.

### 2.1 Search strategy

Human and animal studies were retrieved through multiple electronic journal databases – Medline, Web of Science, EMBASE, Zoological Record, and PsycINFO (for detailed search strategy and search strings, refer to supplemental materials A, Appendix 1). Duplicate entries were removed and an initial title/abstract screening was carried out. Studies were removed from further analysis if all three reviewers independently flagged the entry as clearly irrelevant; i.e. the study included no hGC measurements. Subsequently, all papers citing the retained journal entries – identified using Google Scholar – were retrieved, duplicates were removed, and these additional studies were pooled with the original cohort.

### 2.2 Inclusion/exclusion criteria

Only English-language peer-reviewed papers presenting original data were included for further analysis. Papers had to include quantification of GCs in hair from a vertebrate species, used as a measure of physiological or psychological stress. Two designs were admissible: either hGCs were measured in a group of (purportedly) stressed individuals, related to a less stressed control group (or the same subjects sampled under a less stressful condition), or the measurements were related (correlated) to GC measurements in another biological matrix for the same individuals. Other biological matrices, where GCs have previously been validated to be a measure of central HPA-axis functioning/activity, are blood, saliva, urine and feces (for an overview, refer to e.g. Sheriff et al.^27^). Initial full-text screenings were carried out by three reviewers independently; disagreements on whether to include a study were settled in a meeting where a consensus was reached for all studies without a previously unanimous decision.

### 2.3 Quality assessments

Standardized checklists/protocols for quality assessments have been proposed for human^28,29^ and animal^30,31^ studies. To our knowledge, there are no standardized schemes that can be used, unmodified, for both. Instead, we utilized the method guide developed by the Agency for Healthcare Research and Quality^32,33^ as a foundation for our quality assessments, constructing a protocol for assessing external validity/reporting quality and a nine-point risk-of-bias checklist critically assessing internal validity (Supplemental materials A, Appendices 2 and 3).

### 2.4 Data extractions

Information on study design, including subject characteristics, was extracted along with basic data on sample treatment and analysis from the retained publications. For studies of a stress/control design, the number of subjects was extracted along with means and standard deviations (recalculated from other measures of dispersion, where needed) for the groups. For correlational studies, the number of subjects were extracted together with a correlation coefficient. For studies where only the test of significance was presented, correlation coefficients were calculated using the p-value, where this was reported as an exact number (as opposed to a range, e.g. p < 0.05).

Where values were only reported graphically, data was estimated using on-screen measuring software (Universal Desktop Ruler, AVPSoft). Where data had been transformed to conform to a normal distribution, the transformed values were extracted. If the measure of dispersion was unclear, we assumed the reported values were SEM, thereby producing a conservative estimate. Data were extracted by a non-blinded reviewer, checked independently by a second reviewer, and brought to a three-reviewer consensus meeting if the reviewers disagreed. If data was not extractable, or partially missing, the corresponding author for the study was contacted by email. Where a response could not be obtained after a reminder email, other methods were employed, including (but not limited to) contacting the first/last author of the publication, using ORCID to obtain up-to-date contact details for the corresponding author, and reaching out to authors over ResearchGate. If the missing data could not be produced, or if an author did not respond despite multiple contact attempts over the course of one month, the study in question was removed.

### 2.5 Data analysis

Due to the heterogeneous data material, with absolute concentrations known to differ significantly depending on the analysis method employed^34,35^, and differing preferred ways of reporting (e.g. some authors preferring to report the resulting concentration of the extraction medium, as opposed to the hair content of glucocorticoids), standardized mean differences were employed as the end-point comparison for the stress/control design studies. In order to not artificially inflate the weight of studies with multiple comparisons, best practices for combining study data were employed (refer to Supplemental Materials B): Stressor and control groups were combined in accordance with the methods recommended by the Cochrane Collaboration^36^. Where multiple measurements were obtained from the same individual, these nonindependent measurements were combined utilizing the methods outlined by Borenstein et al.^37^. For repeated measure designs – where multiple hair samples were collected over time, or produced through segmentation of hairs – adjacent samples (e.g. neighboring time-points/hair segments) were assumed to have an average correlation of 0.75 (samples one-over thus correlated by 0.75^2^, etc.) unless a correlation was explicitly stated in the paper. The estimate was based on raw data obtained from the analyzed studies (e.g. Schalinski et al.^38^) and robustness analysis^37^.

Due to the highly heterogeneous study designs and data material, the studies needed to be stratified to distinguish between types of stressor for a meaningful interpretation of data. In the original protocol, comparisons were to be stratified by duration/temporality of the stressor (acute/intermittent/chronic), with PTSD studies analyzed separately; however, timing of the stressor (with respect to the subsequent sampling) proved hard to pin down with exactitude from the reporting. Instead of combining potentially incompatible study designs, a more granular subdivision was employed. Induced (acute) stressors, chronic (non-acute) stressors, and studies of PTSD were separated. However, past stressors – where the subjects were no longer exposed to the stressor at the time of sampling – were separated out from the chronic stress studies (as suggested by the findings of Stalder et al.^26^). Moreover, opportunistically observed stress and self-assessed stress were separated out into their own categories, as the timing and duration of these types of stress was hard to categorize with respect to the time of sampling. Despite the subdivision of data by study design, the stress models within each category were considered diverse enough to where data were synthesized using random effects models.

Correlation coefficients were synthesized by first transforming data onto Fisher’s z scale, according to the method of Hedges and Olkin^39^. Random effects models were employed for each compartment separately, and where multiple coefficients were extracted from a single study, the weights of these z values were adjusted to avoid inflating the weight of any one study. For ease of interpretation, back-transformed data are presented.

Originally, in the study protocol, funnel plot analysis and Egger regressions were suggested as a method for testing for publication bias. The highly heterogeneous data, which had to be stratified into subgroups to facilitate meaningful interpretation, was poorly suited for funnel plot analyses^40^ however. Moreover it has been suggested that the use of standardized mean differences can lead to funnel plot asymmetry even when no publication bias exists^41^. Instead of constructing funnel plots, leave-one-out analysis^42^ was employed to test whether the results of the meta-analyses could be considered robust or whether the overall conclusions could easily be influenced by moderate levels of publication bias.

## 3. Results

**Figure 1.**
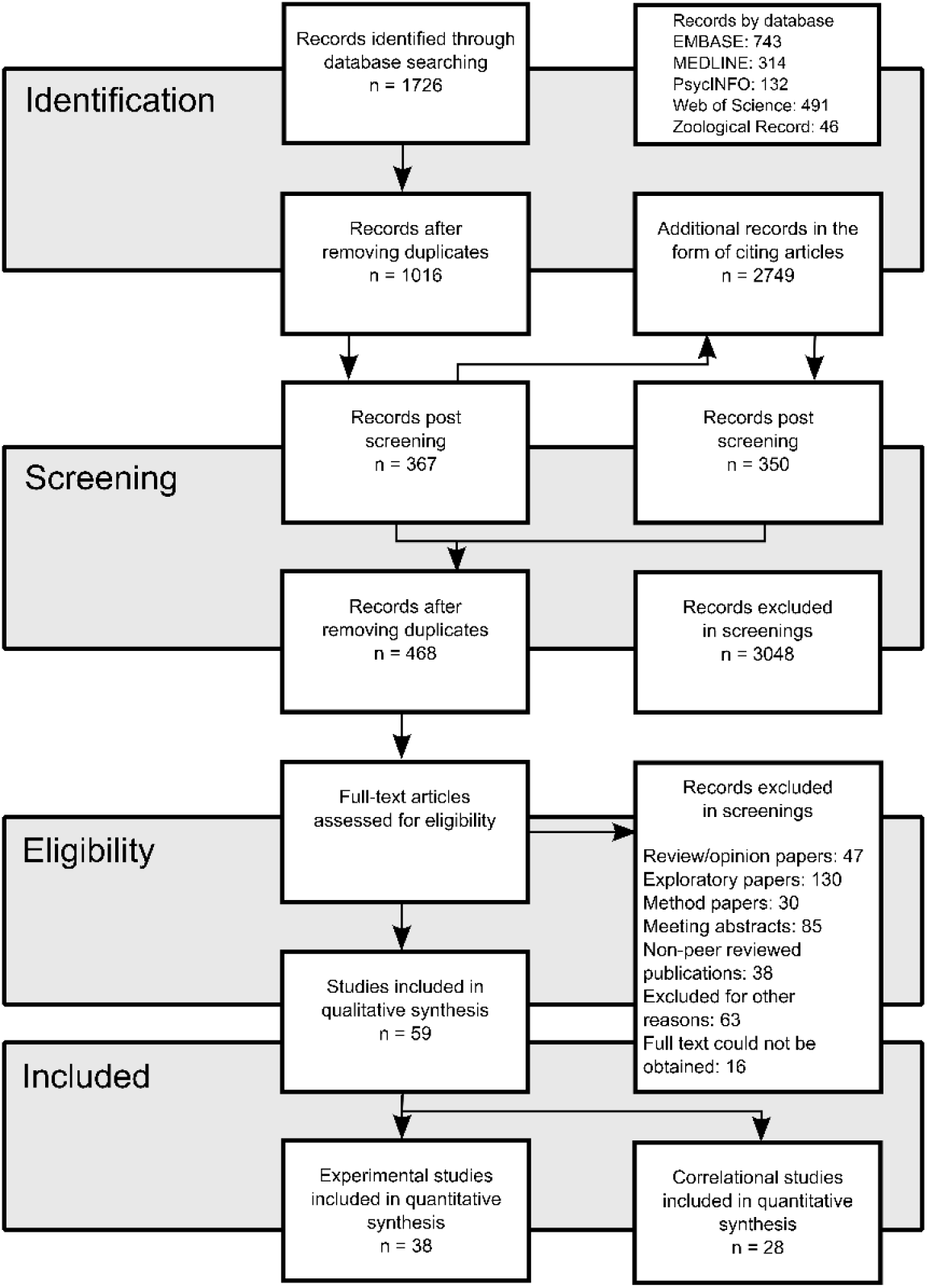
Flow chart outlining the systematic search strategy, the subsequent screening, and inclusion/exclusion of database entries. The diagram has been adapted from the PRISMA Flow Diagram^43^.

A total of 3,518 unique entries were found using the search strategy, where 468 entries were retained for full text analysis. A majority of these studies were subsequently excluded due to not meeting the pre-stated inclusion criteria (Figure 2): 28% were excluded due to their exploratory study design – often characterized by the lack of a control group and a clear a priori hypothesis; 16% presented no data from a controlled study – these were mostly method papers, reviews, opinion papers, and other narrative journal entries; 26% were not peer-reviewed publications – these were mostly meeting abstracts and theses. Other incompatible study designs, and entries where the full text could not be obtained, made up 17% of the entries retained for full text screenings.

**Figure 2.**
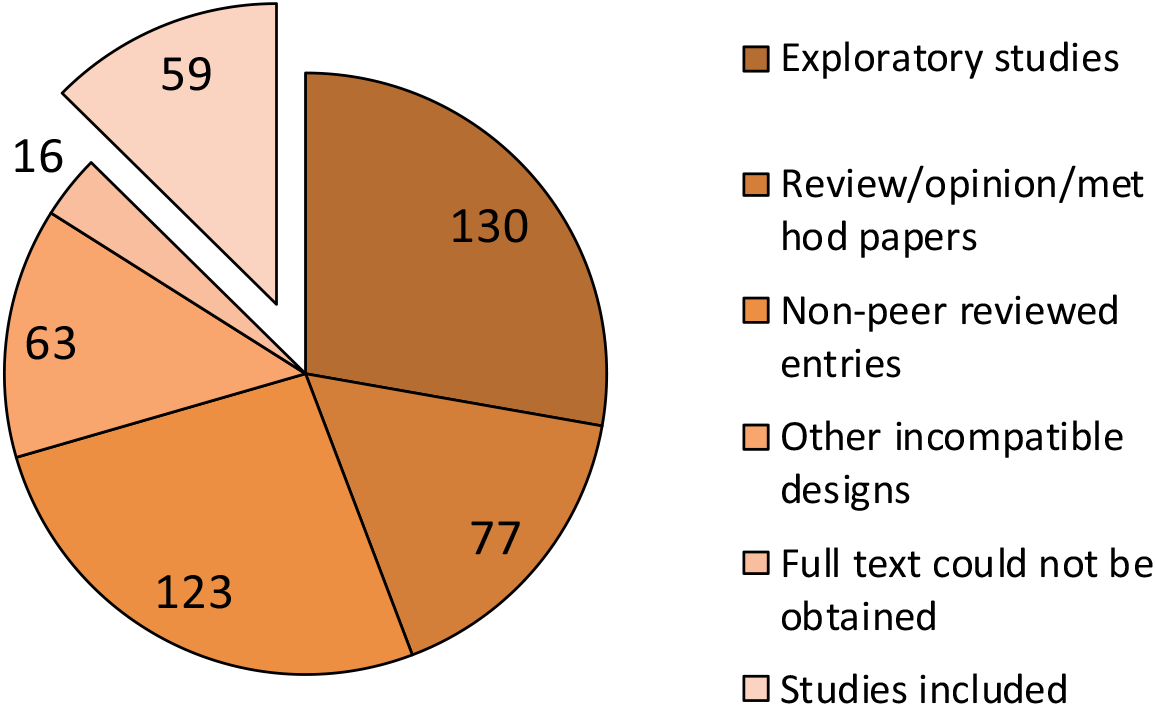
Breakdown of the entries subjected to full-text screening. Notably, less than one in seven publications utilizing/discussing hGCs as a measure of stress could be considered controlled, peer-reviewed, studies contributing empirical evidence.

For the entries retained for full text screening – where all texts were verified to concern the use of hGC – an exponential growth in method adoption is obvious: 2015 saw more publications on hGC than had been published between 2003 and 2011 in total. Presently, a new publication on hGC is available online every three days.

**Figure 3.**
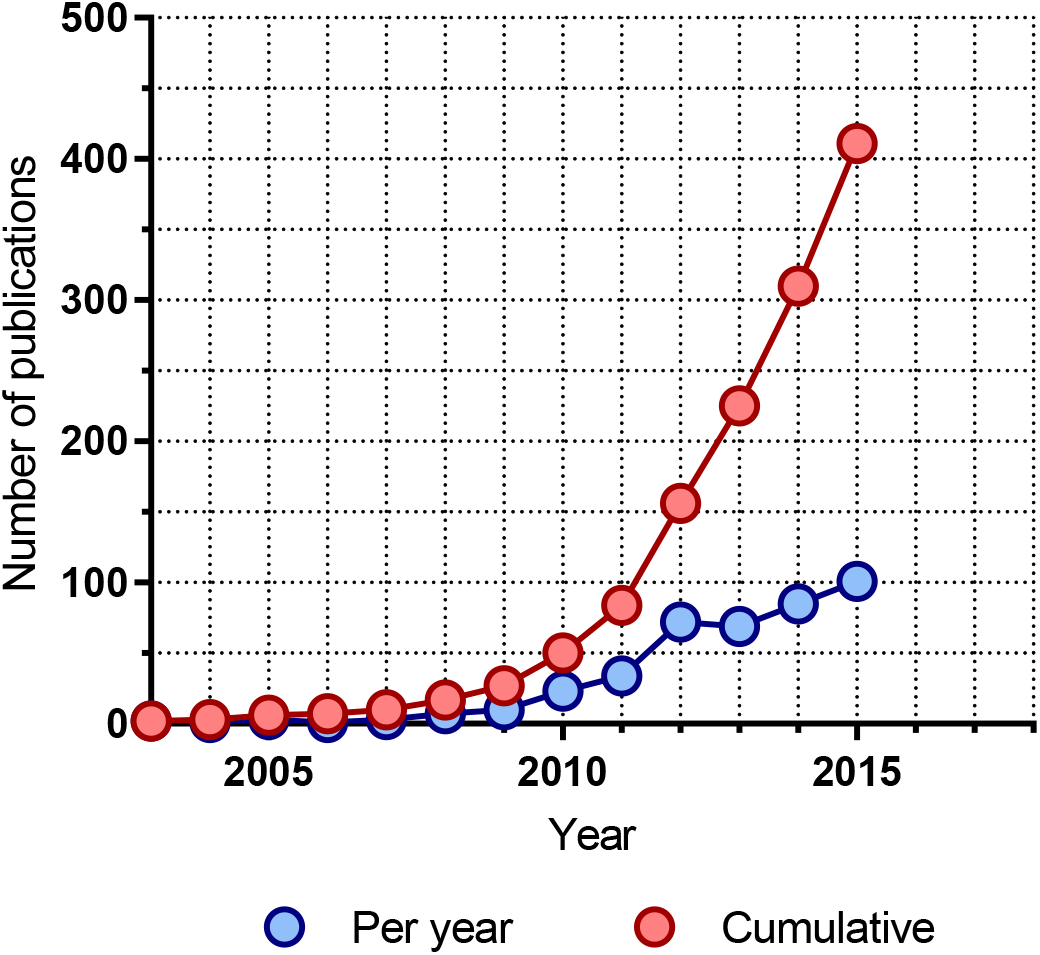
Publication trend 2003-2015 for publications discussing the use of hair glucocorticoids as a measure of stress. The data used to create the graph stem from the 468 unique records identified through the screening process (entries from 2016 have been omitted).

### 3.1 Study quality

Of the 59 peer-reviewed publications included in the present systematic review, 38 papers reported on 43 studies with a stress group/control group design that could be assessed for study quality.

A salient trend was found when assessing the risk of bias: A majority of the 38 papers did not account for the possibility that a stress response other than the one that was purportedly studied could have influenced the results (Figure 4). The influence of concurrent interventions or unintended exposures could only be ruled out in 9 (24%) of the studies. The influence of confounding factors could only be ruled out in 16 (42%) of the studies, and only 12 (32%) of the studies featured a study design that ensured that the subjects were equally exposed to any confounding factors. Similar ambient conditions for stress and control groups could also only be guaranteed in 15 (39%) of the studies. Remarkably, only 3 (8%) of the studies reported on blinding of the outcome assessors, even though this is an explicit recommendation of most present-day best-practice frameworks (e.g. the ARRIVE guidelines^30^). In no one study were all of the sources of bias addressed, and in a few none were (for a by-entry summary of the risk-of-bias analysis, refer to Supplemental materials C, appendix 1).

**Figure 4.**
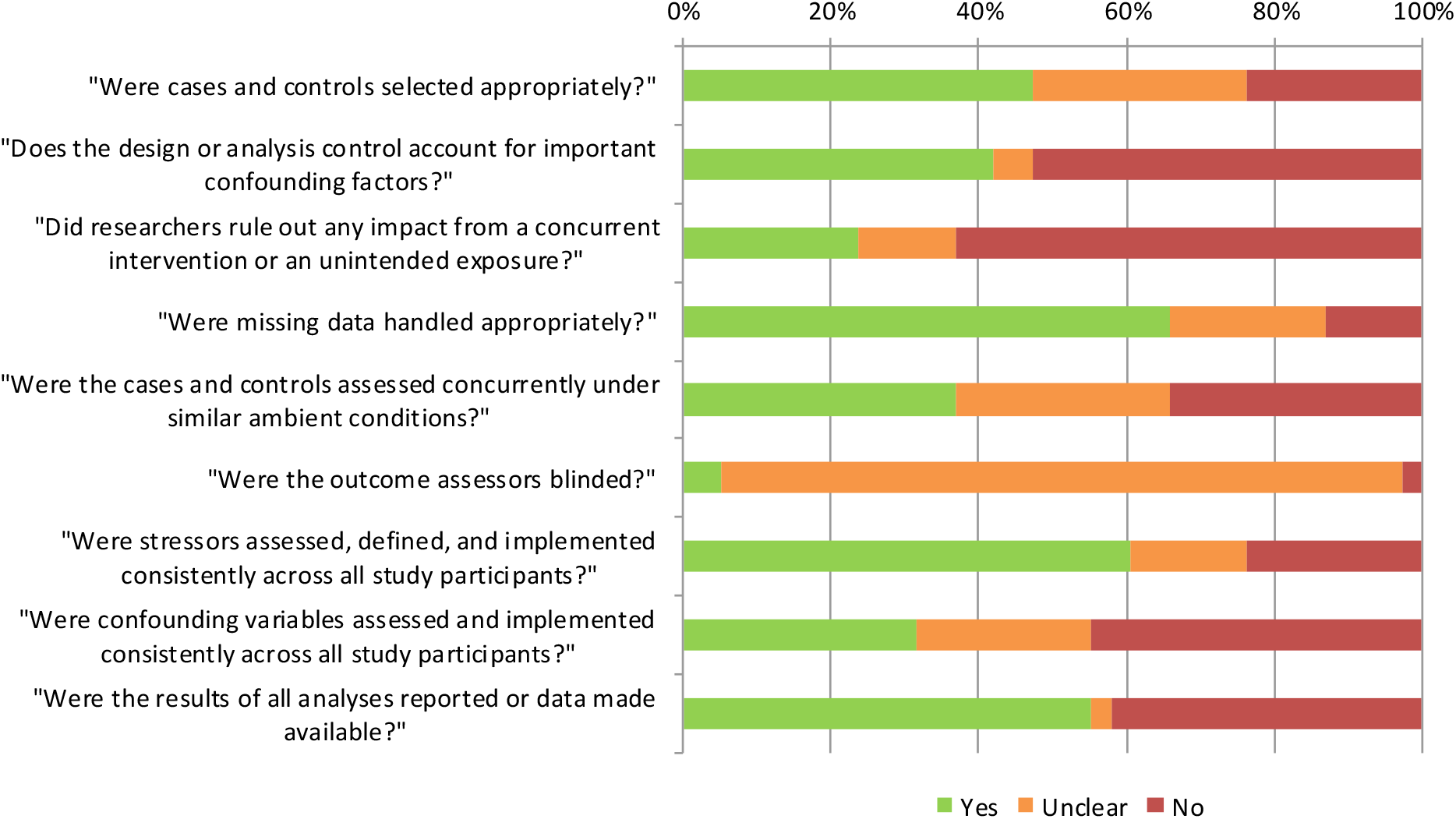
Results from the risk-of-bias checklist assessment of the experimental study designs.

### 3.2 Study characteristics and data extraction

The studies retained for analysis presented a diverse set, with no two study designs quite alike (Tables 1 and 2). Of the studies retained for analysis, roughly half (48%) were human studies. Both sexes have been studied in roughly equal numbers (52% female subjects across all studies), but only rarely were equal sex ratios employed in any one study; study objectives tending to bias the sex ratio in favor of one or the other. We made initial attempts at exploring sex differences – similar to a previous meta-analysis^26^ – however the data was insufficient to draw any conclusions. Human studies were consistent in sampling the posterior vertex of the head, whereas the non-human studies appeared to sample regions by convenience or just by random (e.g. studies in dogs have sampled backs, shoulders, chests and legs, depending on research group and study). Although often discussed as a potential issue^44,45^ no one study admitted to including hair follicles in their hair samples and all but six papers^5,22,46–49^ explicitly described methods designed to ensure samples being free of follicles. Only human and other primate studies employed the “stress calendar” idea, sub-sectioning hairs to infer circulating GC levels at multiple time points in the past from the same sample. A clear majority (54 studies, 84%) of the studies employed a washing step, intended to remove contaminants from the outside of the hairs, and all but two studies minced/pulverized the hairs prior to analysis. For quantification of GCs, antibody-based methods were most frequently employed (53 studies, 83%), however numerous different protocols/antibodies have been utilized.

**Table 1.** Study characteristics for studies with extracted correlations. In total, 30 studies were extracted from 28 peer-reviewed publications, collecting data from 886 subjects across nine species.

| Study | Subjects | Males/females/unknown | Sampling site | Sub-segmentation? | Follicle? | Wash? | Processing? | Analysis method |
| --- | --- | --- | --- | --- | --- | --- | --- | --- |
| Accorsi, 2008 | Cats | 8/19/0 | Back (ischiatric region) | No | No | No | Cut | RIA |
|  | Dogs | 21/8/0 | Back (ischiatric region) | No | No | No | Cut | RIA |
| Bennett, 2010 | Dogs | 0/0/48 | Back (ischiatric region) | No | No | Yes | Milled | ELISA |
| Bryan, 2013a | Dogs | 5/2/0 | Shoulders | No | No | Yes | Milled | ELISA |
| Chan, 2014 | Humans | 26/31/0 | Head | No | No | No | Cut | ELISA |
| Chen, 2014 | Humans | 29/0/0 | Head (posterior vertex) | No | No | Yes | Milled | HPLC-MS/MS |
| Corradini, 2013 | Dogs | 49/41/0 | Chest (xiphoid region) | No | No | No | Cut | RIA |
| D'Anna-Hernandez, 2011 | Humans | 0/21/0 | Head (posterior vertex) | Yes | No | Yes | Milled | ELISA |
| Davenport, 2006 | Rhesus monkeys | 20/0/0 | Neck (posterior vertex) | Yes | No | Yes | Milled | ELISA |
| Kamps, 2014 | Humans | 10/10/0 | Head (posterior vertex) | Yes | No | Yes | Milled | LC-MS/MS |
| Kuehl, 2015 | Humans | 31/54/0 | Head | Unclear | No | Yes | Unclear | CLIA |
| Manenschijn, 2012 | Humans | 0/0/90 | Head (posterior vertex) | No | No | No | Cut | ELISA |
| Mastromonaco, 2014 | Chipmunks | 0/0/62 | Leg | No | No | Yes | Cut | ELISA |
| Moya, 2013 | Cattle | 12/0/0 | Head, neck, shoulder, hip, tail | No | Unclear | Yes | Milled | ELISA |
| Moya, 2015 | Cattle | 80/0/0 | Tail | No | Unclear | Yes | Milled | ELISA |
| Ouschan, 2013 | Dogs | 4/8/0 | Leg (inside) | No | No | Yes | Cut | ELISA |
| Pulopulos, 2014 | Humans | 12/38/0 | Head (posterior vertex) | No | No | Yes | Milled | Unclear |
| Sauvé, 2007 | Humans | 19/20/0 | Head (posterior vertex) | No | No | No | Cut | ELISA |
| Schalinski, 2015 | Humans | 0/28/0 | Head (posterior vertex) | Yes | No | Yes | Unclear | CLIA |
| Steudte, 2011 | Humans | 0/27/0 | Head (posterior vertex) | Yes | No | Yes | Milled | CLIA |
| Steudte, 2013 | Humans | 6/72/0 | Head (posterior vertex) | Yes | No | Yes | Milled | LC-MS/MS |
| Sumra, 2015 | Humans | 0/31/0 | Head | No* | No | Yes | Ground | ELISA |
| Tallo-Parra, 2015 | Cattle | 0/17/0 | Head | No | No | Yes | Cut | ELISA |
| van Holland, 2012 | Humans | 0/0/27 | Head | No | No | Yes | Milled | CLIA |
| Vanaelst, 2012 | Humans | 0/32/0 | Head (posterior vertex) | No* | No | No | Cut | LC-MS/MS |
| Wippert, 2014 | Humans | 3/10/0 | Head (posterior vertex) | No | No | Yes | Unclear | LC-MS/MS |
| Xie, 2011 | Humans | 32/0/0 | Head (posterior vertex) | No* | No | Yes | Milled | LC-MS/MS |
| Yamanashi, 2013 | Chimpanzees | 9/0/0 | Arm | No | No | Yes | Milled | ELISA |
| Yu, 2015 | Mice | 19/0/0 | Back | No | No | Yes | Cut | LC-MS/MS |
|  | Rats | 22/0/0 | Back | No | No | Yes | Cut | LC-MS/MS |

**Table 2.**
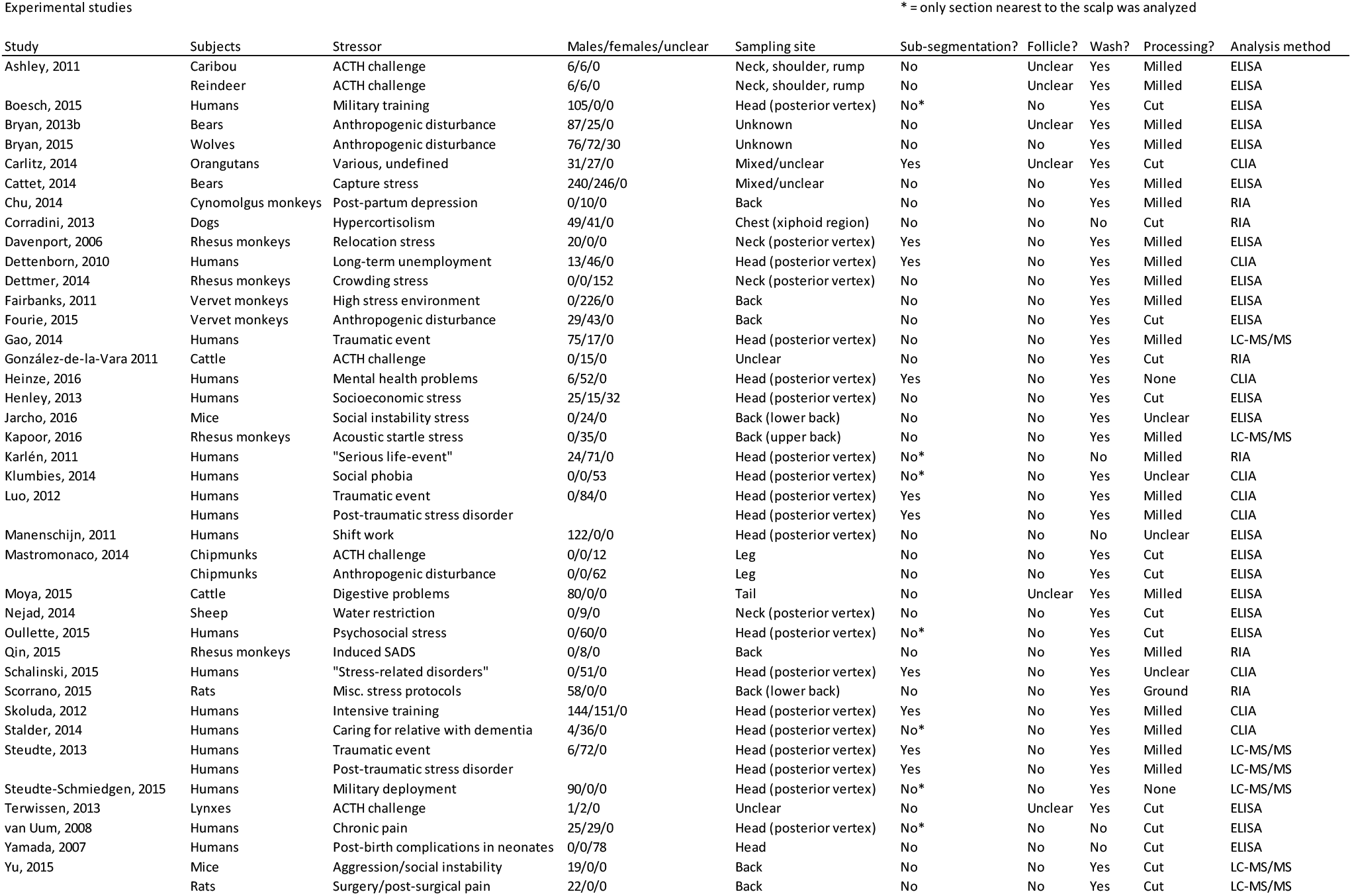
Study characteristics for experimental studies. In total, 43 studies were extracted from 38 peer-reviewed publications, collecting data from 2,842 subjects across 16 species.

When extracting data, two studies – Manenschijn et al.^50^ and Luo et al.^4^ – were singled out as having a reported precision an order of magnitude higher than the other 38 studies (including studies utilizing the very same methodology in comparable subjects). We believe that this is simply due to incorrect reporting of the measure of dispersion. Unable to reach the authors for a comment – despite multiple attempts – we have tentatively included the data from these studies, assuming that the graphically presented measures of dispersion were in fact SEMs, rather than – as listed – 95% CIs.

### 3.3 Correlations with GC in other matrices

Meta-analyses of correlation coefficients revealed a great deal of heterogeneity between studies, as could be expected from the diverse set of studies analyzed (Figure 5). Significant synthesized meta-correlations could be found between hGCs and GCs in blood, saliva and feces. A significant correlation could not be found between hGCs and GCs in urine, however this analysis featured only five studies (collecting 169 subjects), all with fairly high intra-study variance of data. Leave-one-out analysis furthermore revealed that the statistically significant correlation found between GCs in blood and hGCs could not be substantiated if data from the study by Yu et al.^51^ were removed (Table 3).

**Figure 5.**
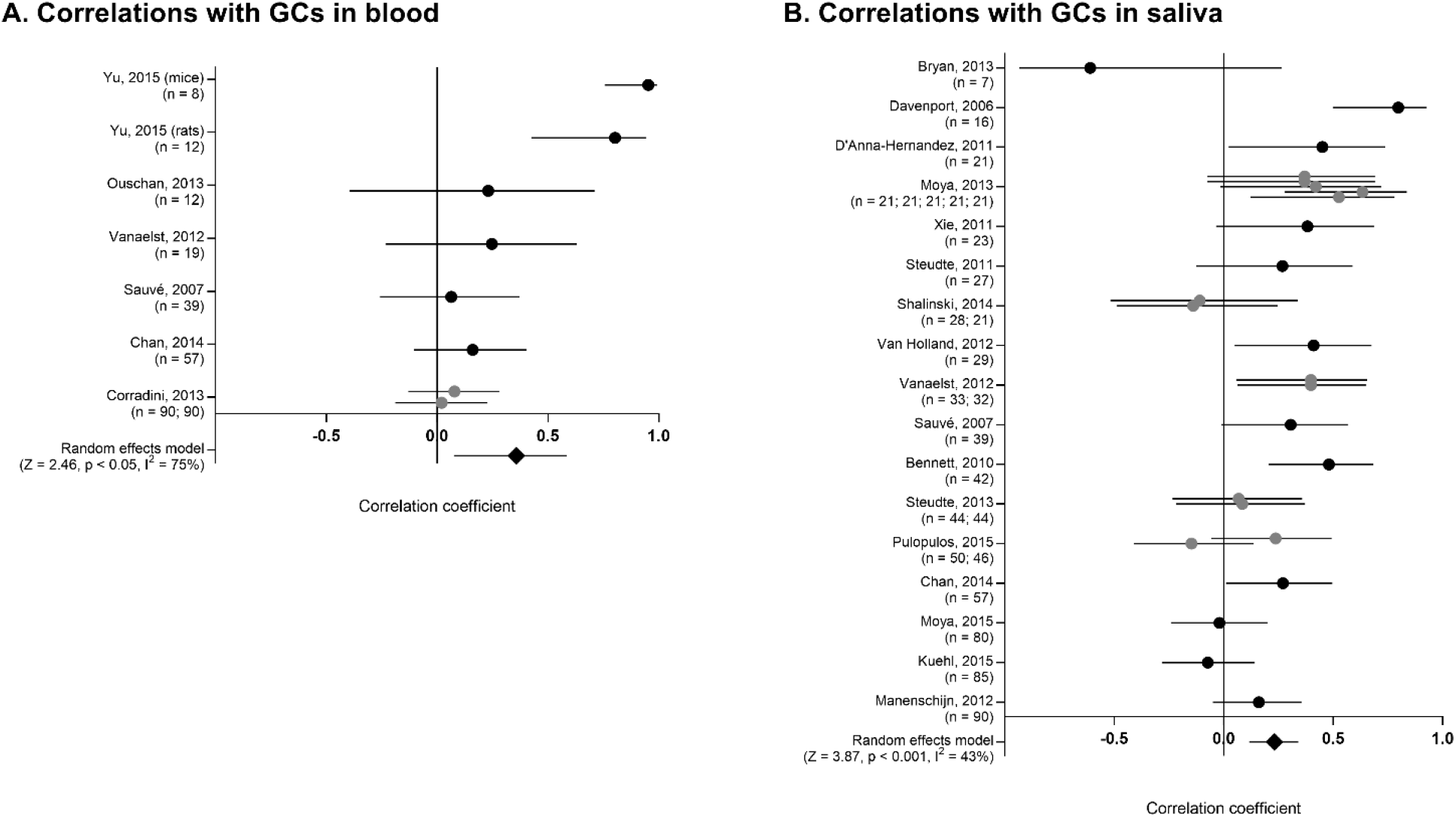
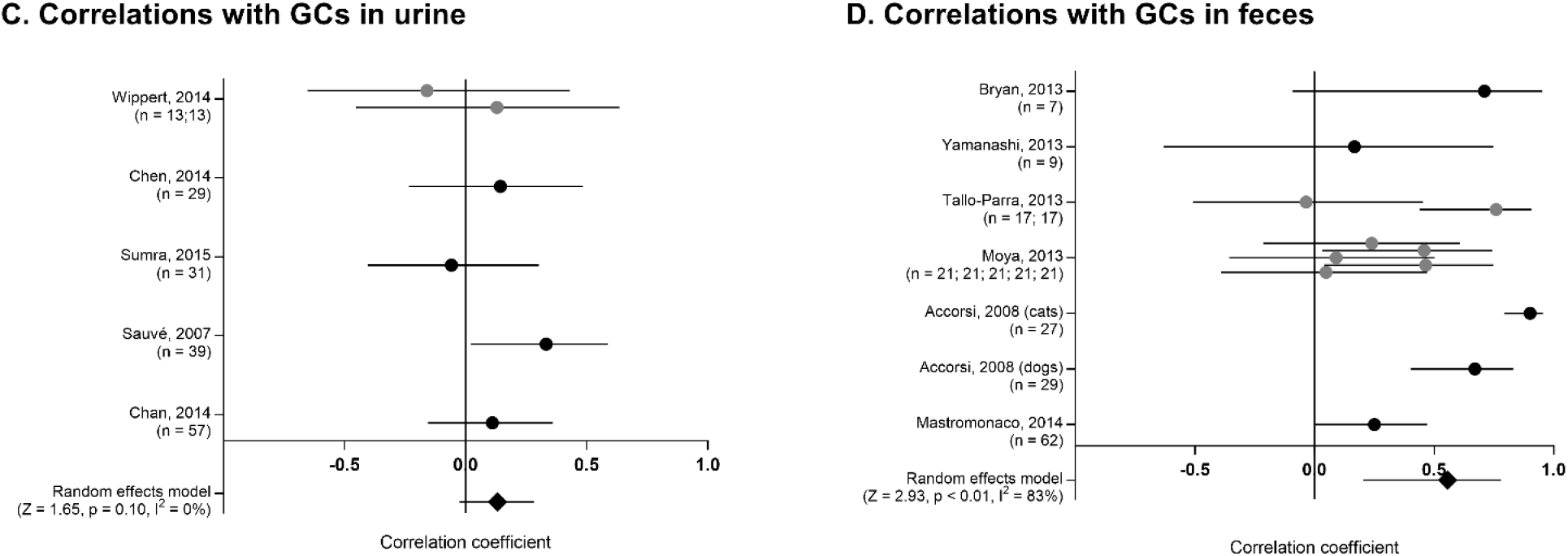
Synthesis of correlation coefficients. Forest plots are presented for correlation coefficients between hGCs and GC in (A) blood, (B) saliva, (C) urine, and (D) feces. Where multiple coefficients were reported in the same study (grey markers) these were used to construct a weighted average for the study before being used in the random effects model.

**Table 3.**
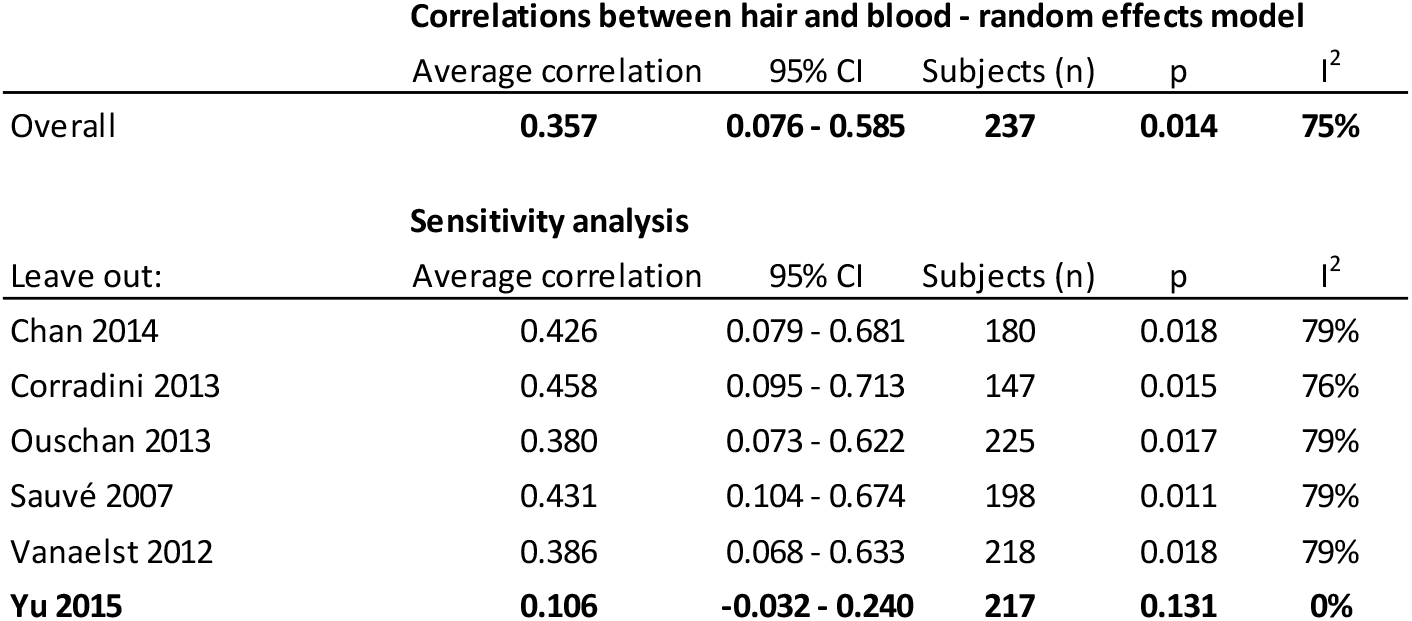
Leave-one-out analysis of correlations between GCs in blood and hair. The greatest difference brought on by removing a single study has been highlighted in bold type. Random effects models have been used throughout.

Correlations between hair and blood - random effects model
|  | Average correlation | 95% CI | Subjects (n) | p | I<sup>2</sup> |
| --- | --- | --- | --- | --- | --- |
| Overall | **0.357** | **0.076 - 0.585** | **237** | **0.014** | **75%** |

| Leave out: | Average correlation | 95% CI | Subjects (n) | p | I<sup>2</sup> |
| --- | --- | --- | --- | --- | --- |
| Chan 2014 | 0.426 | 0.079 - 0.681 | 180 | 0.018 | 79% |
| Corradini 2013 | 0.458 | 0.095 - 0.713 | 147 | 0.015 | 76% |
| Ouschan 2013 | 0.380 | 0.073 - 0.622 | 225 | 0.017 | 79% |
| Sauvé 2007 | 0.431 | 0.104 - 0.674 | 198 | 0.011 | 79% |
| Vanaelst 2012 | 0.386 | 0.068 - 0.633 | 218 | 0.018 | 79% |
| **Yu 2015** | **0.106** | **-0.032 - 0.240** | **217** | **0.131** | **0%** |

Moreover, removing the data from the study by Accorsi et al.^52^ would more than halve the synthesized correlation coefficient between GCs in feces and hGC (putting it in range with the other correlations at r = 0.22), suggesting that the strength of the correlation may be somewhat overestimated (for a complete set of leave-one-out analyses, refer to Supplemental Materials C, Appendix 2).

### 3.4 hGCs as a measure of stress

Induced (acute) stress models produced a clear elevation in GC concentrations measured in hairs (Figure 6) with low inter-study heterogeneity (I^2^ could not be estimated).

**Figure 6.**
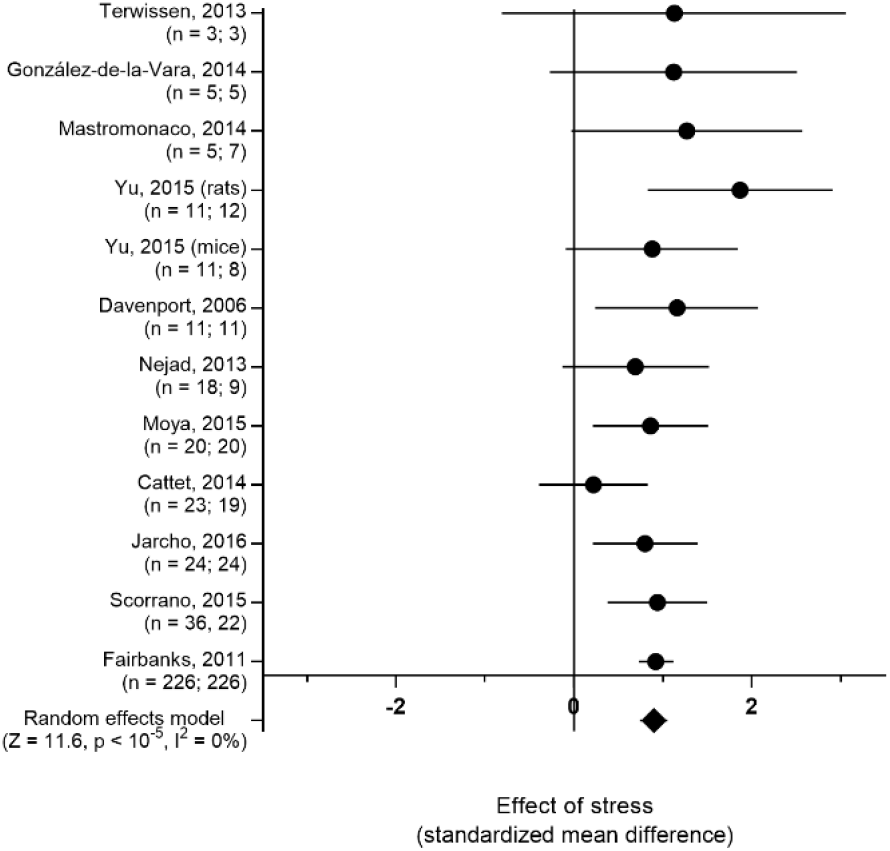
Forest plot summarizing results from induced (acute stress) studies. The number of subjects in the studies are listed with the control group last.

Chronic stressors also produced a significant elevation in deposited GC compared to control groups (Figure 7). The results from the chronic stress studies were however highly heterogeneous suggesting that not all of the studies were comparable. Opportunistically observed stress and self-assessed stress produced unclear results (Figure 8). Finally, stressors that had subsided at the time of sampling did not produce a measurable elevation in hGCs.

**Figure 7.**
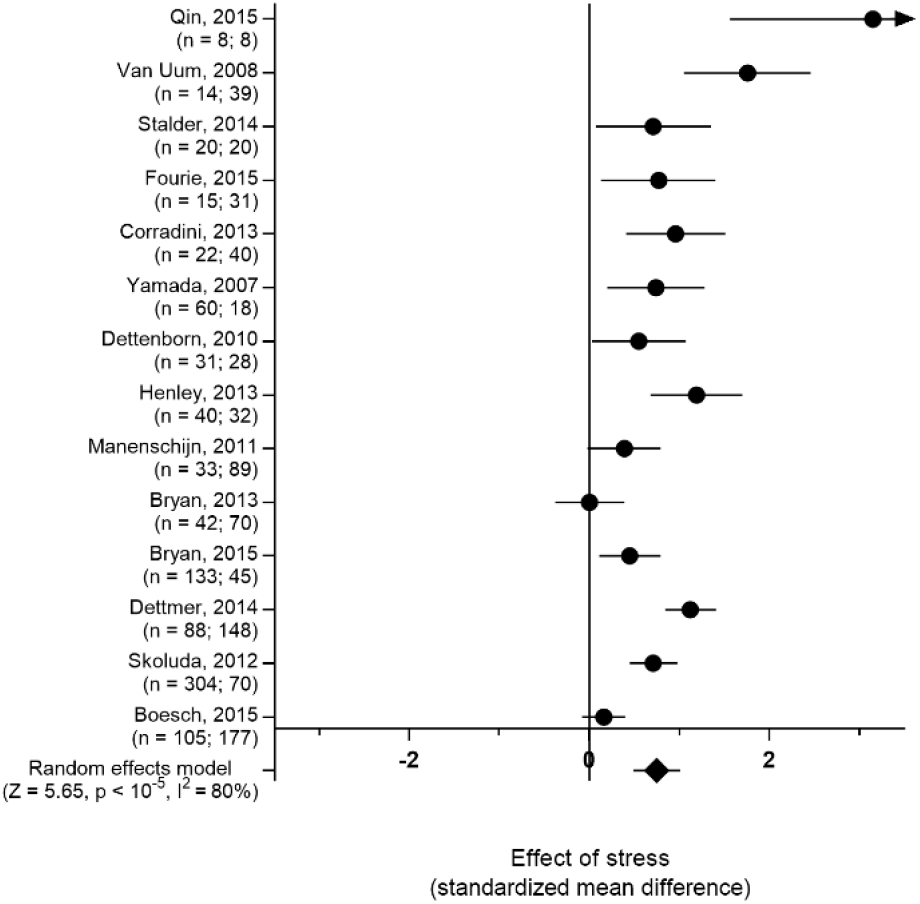
Forest plot summarizing results from chronic stress studies. The number of subjects in the studies are listed with the control group last.

**Figure 8.**
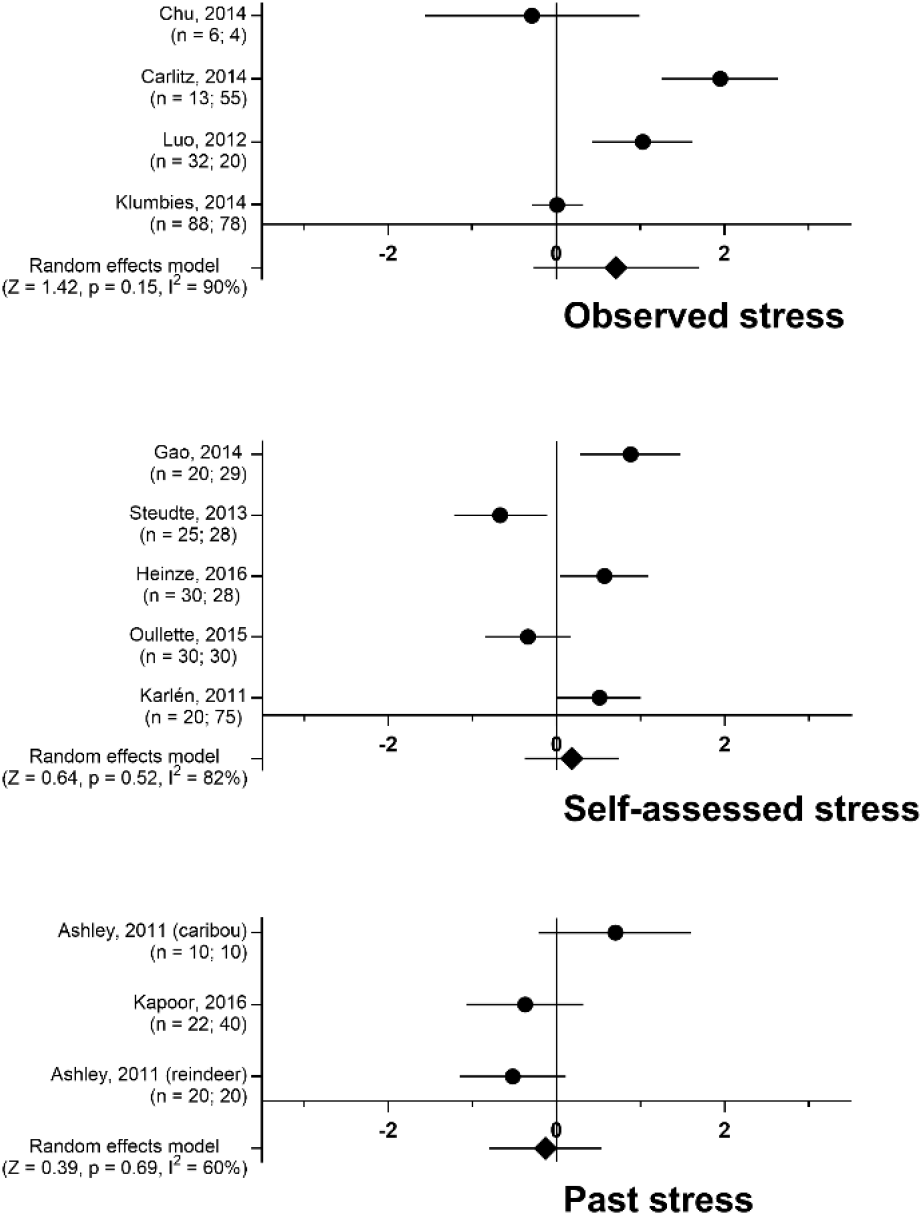
Forest plots summarizing results from studies of inferred (observed/self-assessed stress) and past stress. The number of subjects in the studies are listed with the control group last.

Studies concerning hGCs measured in PTSD sufferers similarly threw up unclear results, with a combination of studies showing both elevations and decreases in hGC output relative to a control group (Figure 9).

**Figure 9.**
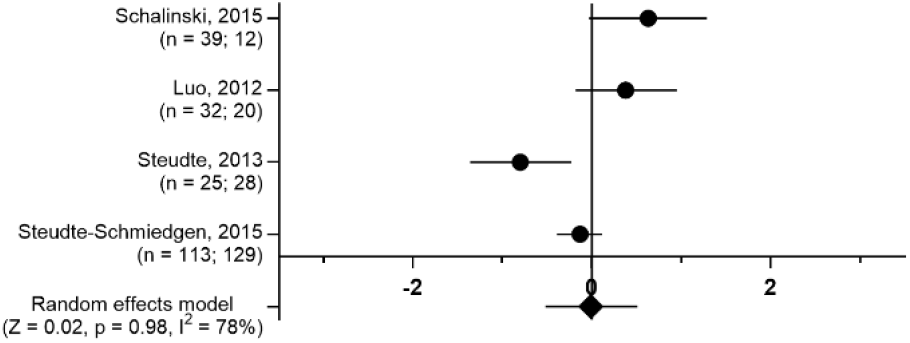
Forest plot summarizing results from studies concerning PTSD. The number of subjects in the studies are listed with the control group last.

## 4. Discussion

With a couple of new papers appearing every week that concern or utilize hGC analysis, it is fair to say that it has become a widespread method for assessing stress. But with hair-growth being a slow process, and popular speculation^53,54^ suggesting that GCs are sequestered by hairs over several weeks – if not months – controlled studies are hard to design and execute. Perhaps this is why our search strategy turned up more narrative reviews, opinion papers and book chapters lauding the method than it did actual controlled studies providing empirical evidence that the method is a sound one. Moreover, the representative study employing hGC analyses, published prior to February 2016, was an exploratory one. Typically, a single cohort of subjects had hair samples collected along with a number of other environmental, physiological, psychological and/or demographic data. Correlations were then constructed to scrutinize which parameters were linked to elevated hGC concentrations. The topics of these studies are varied, from investigations of environmental effects on squirrel gliders^55^ or social effects on German shepherds^56^ to probing cultural^57^, environmental^58^, nutritional^20^ or genetic^59^ influences on psychological stress in people of differing ages. The implicit prior assumption for studies of this kind is that hGCs are linked to central HPA axis functioning and are thus a measure of (chronic) stress. This puts even more of an onus on the (relatively small number of) controlled studies to validate and affirm the use of hGC concentrations as a measure of stress. Even though systematic approaches have been attempted in the past to synthesize data from controlled studies of hGC analyses, a cross-species comparison has, to our knowledge, not been attempted.

The present investigation supports the use of hGCs as a measure of central HPA axis functioning and, consequently, as a stress-sensitive biomarker. The compounded data however calls into question the temporality of the marker, suggesting it is a better marker for ongoing than of past stress.

In studies where subjects were exposed to a controlled stressor, a predictable elevation was found in most cases. Whether repeated ACTH challenges^60^ or a more elaborate protocol combining multiple stressors^61^, a consistent increase was found across species when comparing challenged subjects to unstressed controls.

Whereas most studies sampled hairs at least two weeks after having applied the stressor, the study by Cattet et al.^12^ is remarkable in that they report elevated hGC within hours of stressor onset. In a similar vein, most stress protocols were applied continuously for weeks before hGC concentrations were evaluated, but González-de-la-Vara et al.^60^ found that they could detect an elevation in hGC two weeks after a pair of sustained-release ACTH injections. Notably both highlighted studies employed a washing step in their analyses to ensure that their results were not confounded by external sources of GCs – i.e. GCs in sweat or sebum artificially inflating the hGC measurements. Both studies point to hGC concentrations being reflective, primarily, of events in the recent past, as opposed to historical stressors. This is also consistent with hGCs correlating with GCs in other matrices.

Although both inter- and intra-study variances were high for the collated data, it is clear that hGC concentrations correlate significantly with GC concentrations in other matrices. The synthesized correlation coefficients are weak to moderate – ranging from 0.13 to 0.56 – but this is in range with the correlations between established matrices obtained in these very same studies^62–65^. Due to large fluctuations stemming from the pulsatile nature of GC release^66^, coupled with the different temporality of the matrices – serum and saliva concentrations of GCs change in a matter of minutes in response to a stressor, urinary and fecal GCs change over a period of hours^67^ – these correlations will inevitably be moderate at the most. The correlation between hGCs and GCs in feces is the strongest of the four, which is to be expected as fecal samples integrate circulating GC concentrations over a period of several hours. Hairs are similarly suggested to sequester GCs from circulation over a longer time window. In the face of popular claims, it is unlikely that this time window is several weeks long, however, as hGC concentrations also correlate significantly with serum and salivary concentrations of GCs.

When compiling studies of chronic stress, a link between individuals experiencing stress and elevated levels of hGCs was found, albeit a slightly weaker link than for acute stressors. The greater level of heterogeneity of this dataset is probably part because some of the studies were carried out under highly uncontrolled circumstances. With long-term studies featuring subjects – whether human or non-human – in an uncontrolled environment, it is hard to ensure that the studied stressor is the sole and most influential source of stress. It may be that a lack of dietary salmon elicits a physiological stress reaction in grizzly bears, as suggested by Bryan et al.^46^, but it is quite impossible to tell what other factors might influence the life and allostasis of these bears. The confounding factors of this study may well have drowned out the effect the authors were looking for. Similarly, military training is not all long marches and adrenaline-fueled combat training. With no outside verification, Boesch and collaborators’ soldiers undergoing basic training^68^ may not have had a more active HPA axis than e.g. an office worker with an active lifestyle in the period of sampling. This is not to criticize these experiments; rather, this is to highlight the fact that a number of studies into chronic stressors have an exploratory element to them, as the magnitude of the chronic stressor is hard to judge in relation to a host of ambient stressors. Our risk-of-bias assessment singled out unrelated confounding factors as the most common unchecked source of bias. Only 23% of studies could account for external confounding factors in the studied period, and only in 31% of studies could they be assumed to have been distributed equally between the studied subject groups. The evidence supplied by the chronic stress studies should thus be interpreted carefully.

An important factor shared between the chronic stress studies that demonstrate a clear difference between stressed and control subjects is that the stressor persisted at the time of sampling. When singling out the studies where the stressor could be positively ensured to have subsided at the time of hair sampling, the pattern was found to be different. In the study by Kapoor et al.^69^, pregnant rhesus monkeys were exposed to a daily acoustic startle stress protocol for five weeks. Serum samples analyzed for circulating GC levels were used to verify that the protocol elicited a significant stress response throughout the period. When analyzing hGC concentrations 3-13 weeks later (depending on subject), no elevation could be found relative to a control group; not a trace to be found of a considerable elevation of circulating GC levels persisting for five weeks. Similarly, when Ashley et al. analyzed hairs from both reindeer and caribou two weeks after a single (non-sustained-release) ACTH challenge, no elevation could be found. Fecal GC analyses confirmed that the stressor had subsided after 24-48 hours. With only two studies in this category, we should be careful not to overinterpret; however, this is all part of a recurring pattern. In a recent meta-analysis, Stalder et al.^26^ reanalyzed historical data from human studies in aggregate – collecting data from 66 studies and more than 10,000 hGC samples – and found that in cases of past/absent stress, no relation with elevated hGC concentrations could be found. The idea of hairs containing a historical record of past stress is, and remains, completely unproven, empirical evidence instead pointing to hGCs being a measure of concurrent stress.

In studies where periods of stress were inferred we found the data to be highly heterogeneous. It has been shown before that when human subjects are asked to introspectively assess their own level of stress, assessments correlate poorly with their actual HPA axis functioning^26,70,71^. In the present investigation we see a similar trend for studies relying on observed stress. Whereas we will note that the present investigation contains only a handful of studies, no consistent trend or even weak effect can be inferred. This is not to say that the subjects were not experiencing psychological stress – the studies collect data from distressed subjects ranging from survivors of natural disasters^72^ to patients sourced from mental health services^73,74^ – but it serves as a reminder that the human concept of stress is not synonymous with the prototypical fight-and-flight response. Different states of stress will involve the HPA-axis differently. This is further exemplified by the studies of PTSD, where in two studies^74,75^ a notable reduction in hGCs is found for PTSD subjects when compared to healthy controls. We specifically analyzed PTSD studies separately as it has been suggested that PTSD is accompanied by a lowering in circulating GC levels, as opposed to an elevation. Notably, PTSD subjects are identified through clinical scores suggesting the subjects were assigned to groups according to arbitrary cutoffs in a continuum of chronic stress diagnoses. This muddying of the waters, where the line between chronic stress conditions and PTSD is blurred, may be part in why no clear trend is found concerning hGC profiles for either. In the future, a larger dataset that would allow for a more stringent subgrouping of chronic stress studies based on e.g. clinical scores may assist in identifying more uniform profiles.

Studies where the stressed subjects were identified through observation did similarly not paint a consistent picture. Whether the result of studying animal behavior^76,77^ or of putting human patients through structured interviews^4,78^, there seems to be a mismatch between subjectively assessed stress and hGC levels. For this category of studies we will note that it is particularly concerning that no blinding was employed, even though the findings hinge completely on the subjective assessment of an external observer. Would the marked difference between groups have been as profound in the study by Carlitz et al.^5^ if periods of stress had been determined by a blinded observer and if the stressors had been clearly defined (the study remarkably omits defining the studied stressors)? With few studies and heterogeneous results it is currently hard to determine whether studies that rely on externally assessing stress can provide empirical evidence with respect to the utility of hGC analyses.

Evidence that hGC concentrations are a historical record of stress could also come in the form of studies sub-sectioning hairs, inferring circulating GC levels at multiple time-points in the past. However, the GC levels of hairs were found to be similar across all the sampled segments – individuals with elevated levels of hGCs would have higher levels of hGCs in all segments when compared to controls^73,79,80^. In only two studies the authors attempted to construct a narrative based on point-to-point fluctuations in GC concentrations along the hair shafts. The findings by Luo and collaborators are however marred by a strong wash-out effect, with hGC levels successively becoming lower the further away from the scalp a segment is sourced^4^. The most distant segment is purported to contain the lowest levels of hGCs as this segment is hypothesized to correspond to a period before a major trauma. However, this also holds true for the non-traumatized controls, undermining the hypothesis. The study by Carlitz et al. is similarly problematic in that the narrative seems to have been constructed *post hoc*, and only three individual profiles are shown in the paper^76^. To our knowledge, it is currently all but completely unknown whether the hydrophobic interactions between a steroid hormone and the keratin of a strand of hair are strong enough to lock the molecules permanently in place. Convincing evidence that baleen from whales can trap hormones, leaving a historical record of hormonal fluctuations, has been presented^81,82^. A similar case for hair – a distantly related keratinous matrix – remains elusive however. The difference may lie in the gauge and density of the matrices, with baleen samples being extracted from a depth of several centimeters, using power tools, as opposed to processing the entirety of a micrometer-thick hair. Regardless, the evidence provided by sub-sectioning of hairs, taken altogether, rather seems to suggest that hGCs are distributed evenly across the hair stem. Hydrophobic molecules travelling through the pores formed by a fibrous strand, propelled by capillary forces, is the whole basis for a burning candle where molten wax travels through the wick, defying gravity, ultimately feeding the flame. To assume that a similar effect – longitudinal transport, whether through diffusion or capillary action – could be seen for hydrophobic hormones in hairs does not seem too far-fetched. Reading too much into point-to-point fluctuations thus currently appears to be a case of chasing ghosts.

## 5. Conclusions

The experimental studies using hGC analyses that were carried out prior to February 2016 are spread out across a number of incompatible study designs necessitating sub-group analyses, and consequently diluting the strength of the evidence. Moreover, there is a considerable risk that results from many of the studies have been skewed due to the influence of one, or more, sources of bias. Taken together with the correlational evidence, however, it seems fair to state that hGC levels seem to relate to central HPA axis functioning. GC levels in hairs appear to be an appropriate marker of ongoing physiological stress. If the stressor persists, hGC analyses will remain useful; however, it is currently unadvisable to interpret events in the past based on hGC levels. The idea of GCs being locked into place, providing a historical record of HPA axis functioning has been called into question every time it has been tested in a controlled experiment. Based on the collected evidence we would strongly advice against sub-segmenting hairs, speculating about specific periods in the past. We would be delighted to be proven wrong by a future study, but there is something to be said about, not only the studies our search strategy uncovered, but also the ones that could not be found. Whereas it is hard to design a study where subjects’ stress levels are controlled for weeks on end, it is far from impossible to design a study to test the hypothesis that a stressor in the past can be uncovered in a specific segment of hair. Yet, these studies are nowhere to be found. Whereas the data material did not allow for a stringent exploration of publication bias, it seems highly probable that a number of studies providing negative results have been suppressed. With this review, and others like it, it is our hope that these negative findings may find their way into publications, providing a better picture of when hGC analyses are appropriate, and when they are not.

## Acknowledgements

Academic librarian Kjersti Aksnes-Hopland, University of Bergen Library, assisted in the development of search strategy and search implementation. We would furthermore like to extend our gratitude to the authors who took the time to respond to our queries and supplied missing information.

